# Enterotoxigenic *Escherichia coli* degrades the host MUC2 mucin barrier to facilitate critical pathogen-enterocyte interactions in human small intestine

**DOI:** 10.1101/2021.10.22.465536

**Authors:** Alaullah Sheikh, Tamding Wangdi, Tim J Vickers, Bailey Aaron, Margot Palmer, Mark J. Miller, Seonyoung Kim, Cassandra Herring, Rita Simoes, Jennifer A. Crainic, Jeffrey C. Gildersleeve, Sjoerd van der Post, Gunnar C. Hansson, James M. Fleckenstein

## Abstract

Enterotoxigenic *Escherichia coli* (ETEC) are a genetically diverse pathologic variant of *E. coli* defined by the production of heat-labile (LT) and/or heat-stable (ST) toxins. ETEC are estimated to cause hundreds of millions of cases of diarrheal illness annually. However, it is not clear that all strains are equally equipped to cause disease and asymptomatic colonization with ETEC is common in low-middle income regions lacking basic sanitation and clean water where ETEC are ubiquitous. Recent molecular epidemiology studies have revealed a significant association between strains which produce EatA, a secreted autotransporter protein, and the development of symptomatic infection. Here, we demonstrate that LT stimulates production of MUC2 mucin by goblet cells in human small intestine, enhancing the protective barrier between pathogens and enterocytes. In contrast, using explants of human small intestine as well as small intestinal enteroids, we show that EatA counters this host defense by engaging and degrading the MUC2 mucin barrier to promote bacterial access to target enterocytes and ultimately toxin delivery suggesting that EatA plays a crucial role in the molecular pathogenesis of ETEC. These findings may inform novel approaches to prevention of the acute diarrheal illness as well as the sequelae associated with ETEC and other pathogens that rely on EatA and similar proteases for efficient interaction with their human hosts.

## introduction

Enterotoxigenic *Escherichia coli* (ETEC) are ubiquitous pathogens in low-middle income regions where they are a major cause of morbidity and mortality due to diarrheal illness, particularly among young children(1). In the classical paradigm of ETEC virulence, the organisms adhere to epithelial cells in the small intestine via plasmid-encoded colonization factors where they deliver heat-labile (LT) and/or heat-stable (ST) enterotoxins that promote the net efflux of salt and water into the intestinal lumen with ensuing watery diarrhea. Nevertheless, some features of ETEC illness suggest that this classical paradigm of molecular pathogenesis is far from complete(2). Notably, ETEC cause illness that may range from mild disease to severe diarrhea accompanied by rapid dehydration clinically indistinguishable from cholera(3–6). In addition, ETEC and other bacterial enteric pathogens including *Shigella* have repeatedly been associated with poorly understood non-diarrheal sequelae(7) including environmental enteropathy(8), growth stunting(9–12), malnutrition(13), and cognitive impairment (14, 15).

In addition to the canonical virulence factors, several more recently discovered plasmid-encoded proteins appear to be conserved within the ETEC pathovar(16, 17). Among these is the EatA autotransporter protein(18), a member of the serine protease autotransporters of the *Enterobacteriaceae* (SPATE) family(19). EatA, which shares approximately 74% identity with the SepA autotransporter from *Shigella flexneri(20)*, has recently been shown to degrade MUC2(21), the major gel-forming mucin secreted by goblet cells in both the small and large intestine(22). The 110 kD secreted passenger domain of the EatA autotransporter contains the functional serine protease activity centered on a catalytic triad formed by residues H134, D162, S267(18).

MUC2 is a large (~5200 amino acids) heavily glycosylated protein with more than 80% of its mass comprised of glycans. The central portion of the protein is organized into PTS domains that are comprised largely of repeated proline (P), threonine (T), and serine (S) residues where T and S hydroxyl groups serve as sites for *O*-glycosylation. These densely glycan rich PTS core regions of MUC2 are protected from proteolytic degradation. The MUC2 apoprotein undergoes end-to-end dimerization at the C terminal end of the molecule in the endoplasmic reticulum(23), followed by *O*-glycosylation in the Golgi apparatus, and further multimerization via von Willebrand D domains (VWD) in the N-terminal region of the glycoprotein (24, 25). Ultimately, the MUC2 secreted by goblet cells expands to form large layered polymeric net-like structures that serve as a primary mucosal defense against gastrointestinal pathogens and resident microbiota (26).

Interestingly, recent studies of ETEC isolated from a cohort of young Bangladeshi children followed from birth to two years of age (12) demonstrated that the presence of the *eatA* locus was strongly associated with symptomatic diarrheal disease (27). The studies reported here demonstrate that EatA degradation of mucin plays an essential role in promoting effective pathogen-host interactions central to the molecular pathogenesis of ETEC.

## Results

### ETEC heat-labile toxin stimulates production of a defensive mucin barrier

Although the mucin layer in the small intestine is more penetrable compared to the colon(28), it is possible that it is sufficient to preclude the direct engagement of intestinal epithelial cells ETEC required for effective toxin delivery(29). In addition, studies of rat intestine(30, 31) and cell lines derived from a colonic cancer(32) suggested that cholera toxin, a homologue of LT, can stimulate goblet cell secretion of mucin. Accordingly, LT treatment of enteroids derived from human small intestine resulted in significantly increased transcription of the *MUC2* gene (figure 1a), while treatment of spheroids of small intestinal cells, in which the apical surface of cells is oriented facing an internal luminal cavity, resulted in the release of substantial amounts of MUC2 mucin into the lumen compared to untreated controls (figure 1b). Similarly, treatment of polarized small intestinal enteroid monolayers with LT and cholera toxin, but not a mutant E112K enzymatically inactive version of LT (mLT), resulted in substantial increases in the secretion of MUC2 mucin (figure 1c) and a corresponding increase in integrity (figure 1d). Enteroid monolayers treated with either forskolin or LT, but not untreated monolayers, excluded the migration of fluorescent beads to the epithelial surface suggesting that the induced mucin presents a significant barrier (figure 1e,f). Altogether, these data suggest that both LT and cholera toxin provoke intestinal epithelia to reinforce the protective mucin barrier in the small intestine.

**Figure 1.**
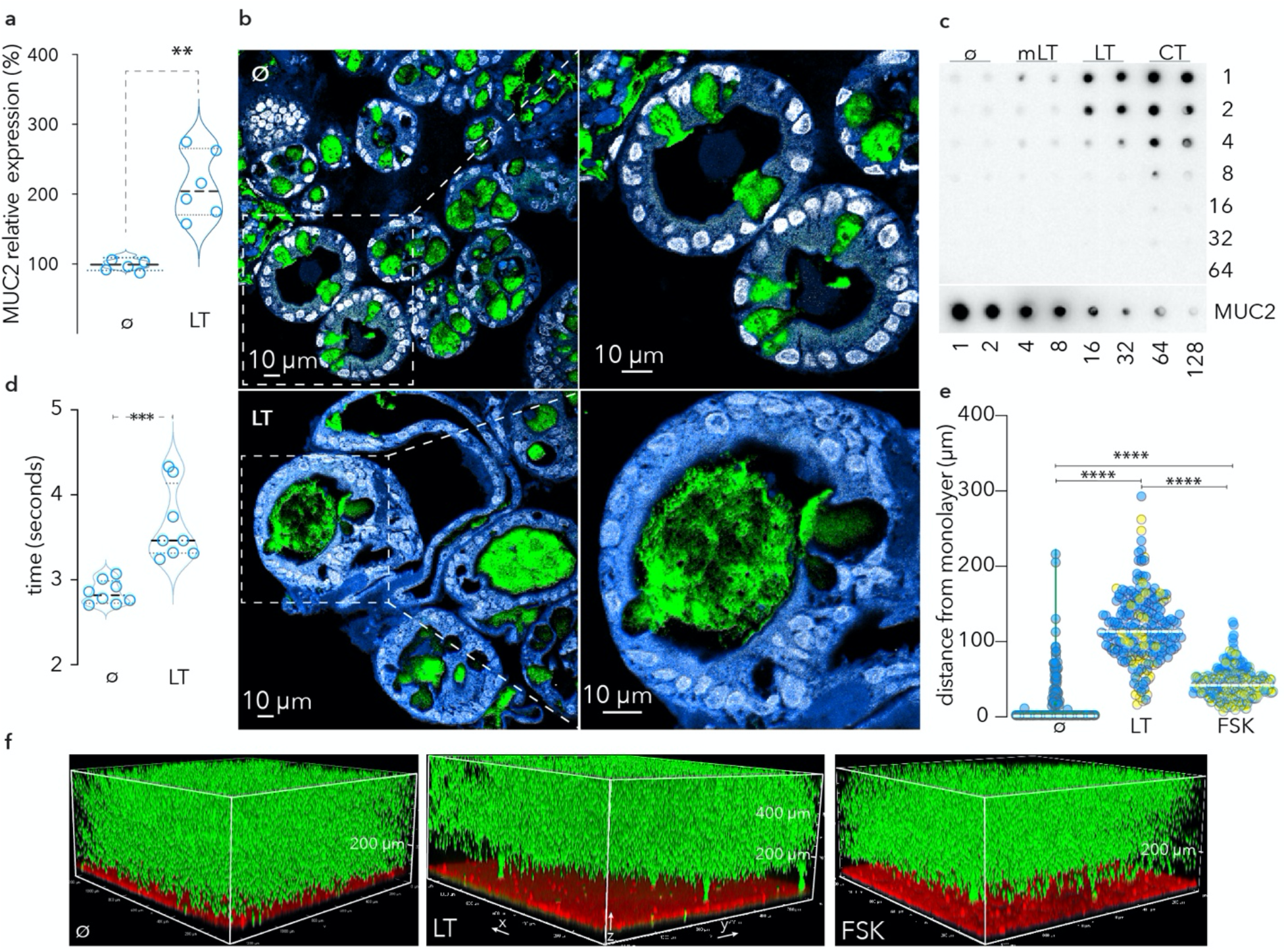
LT stimulates production and secretion of MUC2 mucin barrier by goblet cells in small intestinal epithelia. **a.** transcriptional response (RT-PCR, MUC2) of small intestinal enteroids following treatment with heat-labile toxin (LT, [100 ng/ml] overnight) compared to untreated (ø) control cells. Shown are summary of replicate experiments each with 3 technical replicates (** p=0.02, Mann-Whitney, two-tailed, nonparametric analysis). **b.** small intestinal enteroids (spheroids) in which the apical surface of the enterocytes is oriented to the inside of the sphere. Top panels represent untreated spheroids (ø) where MUC2 (green) is largely contained within goblet cells. Nuclei are pseudo-colored white (DAPI) and cell membranes in blue (CellMask). Hashed line depicts region enlarged in figure below. Bottom panels: LT-treatment of small intestinal spheroids results in MUC2 secretion into the lumen. **c.** MUC2 mucin production by polarized differentiated small intestinal enteroids. α-MUC2 immunoblot shows MUC2 present in apical supernatants of (duplicate wells) untreated monolayers (ø), following treatment with mutant LT (mLT), native LT, or cholera toxin (CT). **d.** LT treatment increases the integrity of the mucin barrier overlying enterocytes. Data represent time required for metal beads to traverse mucin in supernatants obtained from LT-treated or control cells (ø). Shown are results of replicate experiments each with 4 technical replicates (***<0.01 by Mann-Whitney, two-tailed nonparametric testing). **e.** Graph depicts gap distances between 1 μm fluorescent beads and small intestinal enteroid monolayer surfaces comparing LT-treated cells to those treated with forskolin-treated (FSK) or untreated (ø) monolayers. Data from duplicate experiments (n=3 technical replicates) are differentially colored. Each symbol represents the distance (z) between the bead front and the monolayer surface at selected points along the x-y axis. Dashed lines represent median values ****p<0.0001 by Kruskal-Wallis testing. **f.** Volume projections of representative confocal z-stack images of human small intestinal monolayers untreated (ø); treated with LT; or forskolin (FSK). Cells were stained with CellMask (red), treated overnight followed by addition of fluorescent beads (1 μm). Confocal acquisition was ~45 min after addition of beads.

### EatA promotes bacterial migration through the secreted mucin network

To explore the hypothesis that the secreted EatA protease may facilitate pathogen access to host enterocytes by degrading the protective mucin barrier, we first examined the impact of EatA on the viscosity of the mucin matrix. Treatment of purified intestinal mucin with wild type recombinant passenger domain (rEatAp, figure 2a) resulted in significant reduction in the viscosity (figure 2b) relative to either untreated mucin, or mucin treated with proteolytically inactive passenger domain (H134R). Likewise, polyclonal IgG against EatAp prevented reductions in integrity (figure 2c). Next, we found that wild type ETEC migrated significantly faster through purified mucin than the *eatA* mutant (figure 2d), suggesting that mucin serves as a substantial barrier to ETEC migration. Indeed, in *ex vivo* explants of human small intestine (figure 2e, supplemental movie 1), we found that wild type ETEC efficiently penetrated the mucin layer to engage the epithelium, while the *eatA* mutant was excluded from contact with the epithelial surface. In addition, we used small intestinal enteroids in which the *O*-linked MUC2 sugars were metabolically labeled with azide-modified galactosamine (GalNAz) followed by click conjugation with the alkyne flurophore (DBCO-Cy3) to highlight the secreted mucin (red). Again, we found that the *eatA* mutant was trapped by small intestinal mucin (figure 2f, supplemental movie 2).

**Figure 2.**
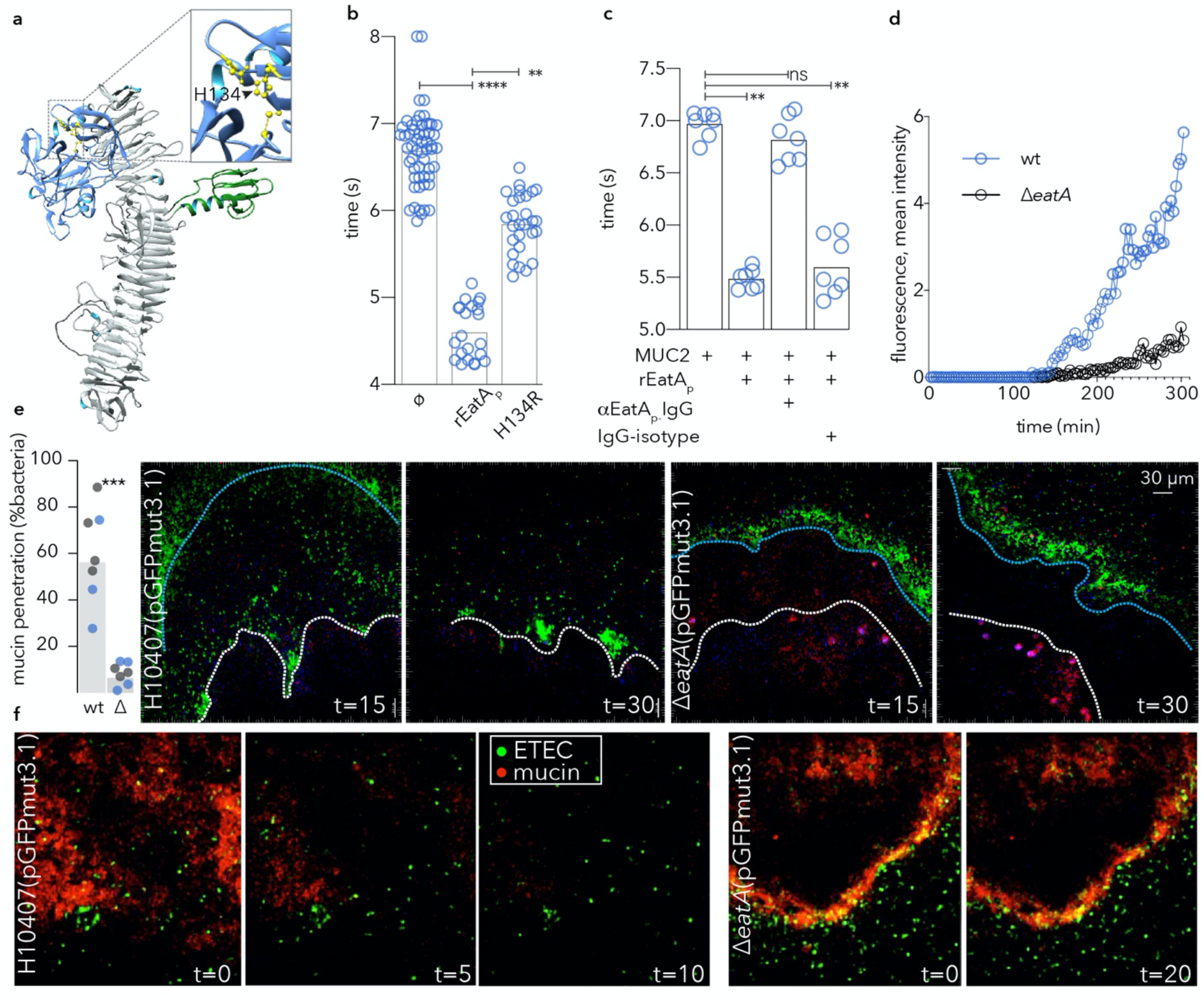
EatA reduces mucin viscosity and promotes bacterial migration through intestinal mucin. **a.** putative EatA passenger domain structure protease subdomain (blue) showing active site with the H134, D162, S267 catalytic triad highlighted in yellow (inset), and the domain of unknown function (DUF) in green. **b.** wild type recombinant EatA passenger domain (rEatA_p_) but not the passenger bearing a mutation in the serine protease catalytic triad (H134R) reduces the integrity of purified MUC2 mucin. Symbols represent technical replicates combined from a total of n=5 experimental replicates. **c.** antibodies against rEatAp inhibit EatA effects on mucin integrity. (Symbols = technical replicates from n=2 experiments). **d.** EatA is required for efficient ETEC migration through purified MUC2 mucin. Each symbol represents average accumulated fluorescence intensity of gfpmut3.1-expressing wild type and *eatA* mutant bacteria over time. Images were acquired ~14 mm from the point of entry. **e.** EatA is required for penetration of the mucin layer in human intestinal explants *in vitro*. Graph at left depicts the % of wild type (wt) and *eatA* mutant (Δ) bacteria expressing pGFPmut3.1 which have penetrated the mucin layer (3 technical replicates in each of two independent experiments from separate tissue donors are shown by color). p=0.0006 by ANOVA. Shown in panels on right are representative images acquired by two photon microscopy of bacteria (green) and penetration into the mucin layer over time. The edge of the intestinal mucin is depicted by the blue line, and the surface of the intestine by the white line. Intestinal cells auto-fluoresce purple**. f.** 2-photon microscopy images of small intestinal enteroids metabolically labeled with GalNAz to highlight secreted mucin (red).

### EatA is required for efficient pathogen-host interaction and toxin delivery

Efficient delivery of ETEC enterotoxin requires intimate interaction of the bacteria with the surface of target intestinal epithelial cells (IECs)(29). We found that in the absence of *eatA* ETEC bacteria were incapable of effectively migrating through MUC2 purified from small intestinal enteroids and were as impaired as an immotile (*fliC*) mutant, while pretreatment of MUC2 with rEatAp restored migration (figure 3a). Similarly, adherence of the *eatA* mutant ETEC to target small intestinal enteroid epithelial cells (figure 3b, supplemental figure 1) was significantly impaired relative to wild type ETEC as was delivery of heat-labile toxin (figure 3c). Collectively, these data suggest that *eatA* plays an essential role in promoting effective ETEC pathogen-host interactions.

**Figure 3.**
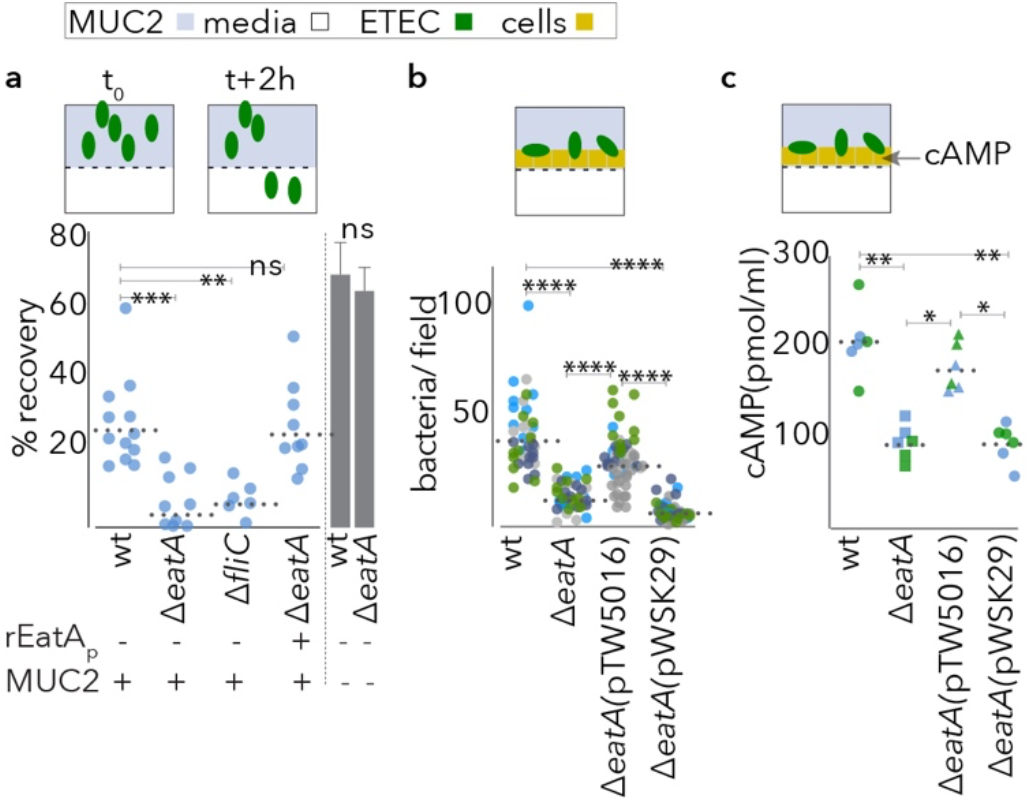
Both EatA and motility are required to penetrate MUC2 and engage small intestinal epithelia. **a.** Shown are data from Transwell mucin migration assays in which wild type H10407(wt), the *eatA*, or flagellin (*fliC*) mutants (~10^5^ cfu) were added to upper chamber containing MUC2 mucin harvested from small intestinal enteroids. Data are expressed as the % of inoculum recovered from the lower chamber after 2 h. rEatA_p_ represents complementation with exogenous EatA passenger domain (50 μg/ml) added at the time of infection. Bar graphs at right demonstrate mean ± SEM recovery in the absence of MUC2**. b.** EatA facilitates bacterial access to epithelial cells. Shown are wt or mutant bacteria adherent to small intestinal enteroid monolayer epithelial cells. The *eatA* mutant was complemented with either a recombinant *eatA* expression plasmid (pTW5016) or the vector control (pWSK29). Shown are the results of three replicate experiments (separated by color) with each symbol (n=40) representing a technical replicate. **c.** EatA is required for efficient ETEC delivery of heat-labile toxin to enteroid monolayers. (shown are cAMP levels in target epithelial cells infected with wild type or mutant bacteria 2 h after infection). P values in each panel reflect comparisons by ANOVA with non-parametric Kruskal-Wallis testing (*p=0.03, **=0.002,***=0.0002, ****<0.0001).

### Identification of a subdomain required for binding and degradation of MUC2 mucin

MUC2 mucin produced by goblet cells in the small intestine is heavily glycosylated. Interestingly, recent structural analysis of SepA suggested that an 80 amino acid subdomain within the passenger domain could represent a carbohydrate binding module(33) (CBM). To investigate the importance of the corresponding region in EatA (figure 4a) to virulence, we first replaced the passenger subdomain of unknown function encompassed by amino acids E541-S616 with a stretch of 8 histidines (DUF∷H_8_). The resulting protein was secreted as efficiently as the parent molecule (figure 4b) and retained the ability to degrade the AAPL synthetic peptide (figure 4c). While the passenger domain containing a single point mutation in the catalytic triad (H134R) colocalized with MUC2 in sections of human intestine (figure 4d), we observed appreciably less interaction between MUC2 and the DUF∷H8 mutant in sections of human ileum (supplemental figure 2, figure 4e) and in the lumen of small intestinal spheroids (figure 4f). Similarly, the DUF∷H_8_ passenger protein exhibited only weak interaction with purified MUC2 (figure 4g), and the mutant protein was incapable of efficiently degrading MUC2 (figure 4h, supplmental figure 3A). Importantly, antibodies specific to the E541-S616 peptide inhibited the activity of the parent molecule (supplemental figure 3B-D). Complementation of the *eatA* mutant *in trans* with a plasmid expressing the DUF∷H8 protein also failed to restore ETEC interaction with target epithelial cells (figure 4i), further suggesting that this region is essential to EatA function. Although interactions of EatA with heavily glycosylated MUC2 molecules may involve carbohydrate moieties, we did not observe significant binding to glycan microarrays (supplemental dataset 2) with either the full length recombinant EatA passenger domain or the mutant lacking the putative carbohydrate binding module encompassed by the E541-S616 subdomain. Likewise, nanoparticles in which Spy-tagged EatA (K535-S616) was conjugated to the surface of SpyCatcher-mi3 (34, 35) were not sufficient to direct MUC2 binding (supplemental figure 4). Collectively, these studies suggest that while this region of the passenger domain is critical for mucin interactions, it likely acts in concert with the proteolytic subdomain to engage and degrade MUC2.

**Figure 4.**
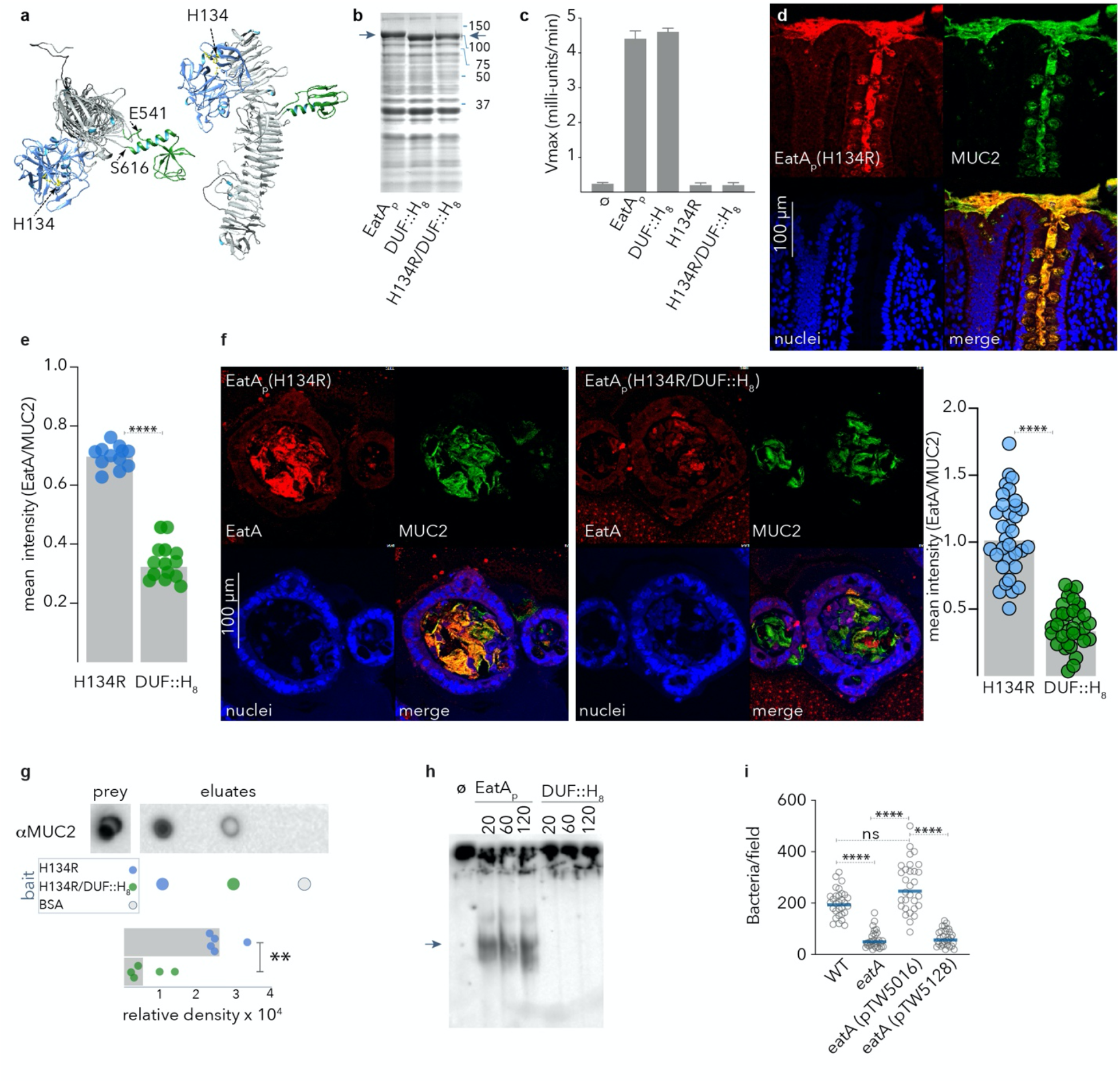
**a.** model of the EatA passenger domain showing the relative location of the protease (blue) and putative CBM (green). **b.** Coomassie-stained SDS_PAGE of TCA-precipitated culture supernatants showing migration of the recombinant passenger domain (arrows) from *E. coli* Ig10B expressing EatA, EatA bearing a polyhistine replacement of the domain of unknown function (DUF∷H_8_), and the H134R/DUF∷H_8_ double mutant passenger protein. **c.** The domain of unknown function is not required for proteolytic activity. Shown are the results of AAPL oligopeptide cleavage experiments for using the wild type EatA passenger protein (EatAp) and mutant passenger proteins. Results represent mean and standard deviation from 3 separate experiments. **d.** EatA passenger domain colocalizes with MUC2 mucin. Panels show representative confocal images of proteolytically inactive EatA (H134R) passenger domain interacting with MUC2 secreted into the lumen of human intestinal explants and retained in goblet cells. **e**. Optimal binding to small intestinal MUC2 requires the subdomain from E541-S616. Graph depicts colocalized binding of H134R or H134R/DUF∷H8 protein to MUC2 in sections of small intestine shown in supplemental figure 2. **f.** Binding to secreted MUC2 requires the E541-S616 region. Shown are maximal intensity projections of confocal microscopy z-stack images of sections of small intestinal spheroids stimulated with heat labile toxin to stimulate release of MUC2 into the lumen. Replicate daughter sections were incubated with the H134R passenger protein (left) or the DUF∷H_8_ mutant protein (right). Graph depicts binding of EatA H134R and DUF∷H_8_ passenger domains relative to target MUC2. (n=33 regions from 30 spheroids; ****<0.0001, two-tailed Mann Whitney). **g.** MUC2 pull-down using EatA(H134R) passenger domain or the H134R/DUF∷H_8_ passenger domain as bait. Graph represents summary of three replicate experiments (**p=0.008) in two-tailed Mann Whitney nonparametric testing. **h.** MUC2 degradation requires the domain of unknown function **i.** optimal ETEC engagement of small intestinal enteroids requires the E541-Q617 region of the passenger domain. Shown are the combined results of three experimental replicates each with 10 technical replicates/experiment (n=30/group). ANOVA (Kruskal-Wallis)****<0.0001..

## Discussion

The studies presented here potentially provide a biological basis for the recent finding that the presence of *eatA* is strongly associated with development of symptomatic disease in young children infected with ETEC(27) and suggest that mucin degradation plays a critical role for these pathogens. It is clear from earlier studies that enterotoxin delivery by these bacteria requires direct enterocyte engagement (29), and the data presented here indicate that in the absence of an effective mucin-degrading enzyme, even the relatively loose layer of mucin overlying the small intestinal epithelia appears to be sufficient to mitigate interaction of ETEC with the host.

This barrier is further reinforced by the host response to LT. Using enteroids propagated from human small intestine, we show that LT and the closely related cholera toxin are potent secretagogues for MUC2 mucin. Therefore, while the layer of mucin overlying small intestinal epithelia is thought to be more penetrable relative to that in the colon, it does not appear to be static; and EatA may have evolved to counter what would otherwise be an effective host defense strategy.

These data may also add another layer of complexity to the interpretation of epidemiologic studies that have to date defined the ETEC pathovar simply by the presence of heat-labile and heat-stable toxins. The large proportion of asymptomatic ETEC colonization in these studies potentially confounds attempts to accurately assess the burden of diarrheal illness attributable to ETEC(36). The present studies provide further data in support of the concept that ETEC are not equally equipped to cause disease. Notably, *eatA* originally discovered on a virulence plasmid of H10407(18), a strain isolated from a patient with cholera-like illness in Bangladesh, has been identified subsequently in other strains with clear metadata linking them to severe cholera-like illness(5, 6). H10407 has been the subject of multiple controlled human infection studies, largely because relative to other strains it reliably causes significant diarrheal illness(37–39).

The presence of SepA, a homologue that shares more than 70% identity with EatA, has been strongly linked to virulence of other pathogens, including enteroaggregative *E. coli* (EAEC) *(40, 41)*. Indeed, in the interrogation of more than 90 genomes from EAEC strains collected in the Global Enteric Multicenter Study (GEMS)(1) for potential virulence genes, *sepA* was recently shown to have the strongest association with the development of moderate-to-severe diarrheal illness(41). The role of SepA in the virulence of *Shigella* species, including *Shigella flexneri* 5a, where SepA was originally discovered(20) on the large virulence plasmid, is currently unknown. However, given that some *Shigella flexneri* 2a as well as other *Shigella* species including *Sonnei* make EatA(42), might indicate that these pathogens also utilize these proteases to degrade the thicker layer of mucin in the colon.

Collectively, the information emerging from molecular epidemiology studies(27), as well as the data presented here, suggest that EatA plays a crucial role in the molecular pathogenesis of ETEC. These findings may inform novel approaches to prevention of the acute diarrheal illness as well as the sequelae associated with ETEC and other pathogens that rely on EatA and similar proteases for efficient interaction with human hosts.

## Materials and Methods

### Bacterial strains, plasmids and growth conditions

Bacterial strains used in this study are presented in Table1. Bacterial cultures were routinely incubated with aeration at 37°C in Luria-Bertani (LB) broth (10g tryptone, 5g yeast extract, and 10g NaCl per liter) or on LB agar plates unless indicated otherwise. Antibiotics were added as appropriate. For infection assays, overnight grown liquid cultures diluted to 1:100 in LB broth were incubated for ~2h to mid log phase.

### Propagation of enteroids and epithelial monolayer culture

Detailed procedures for growth and maintenance of these cells have been described previously (43, 44). Briefly, cryopreserved cells were resuspended in Matrigel (BD Biosciences), and 15 μL of the suspension was seeded into each well of a 48-well tissue culture plate. Cells were maintained in 50% conditioned medium (CM) consisting of 1:1 mixture of L-WRN conditioned media and primary tissue culture media- Advanced DMEM/F12 containing 20% fetal bovine serum (FBS), 2 mM L-glutamine, 100 units/mL penicillin and 0.1 mg/mL streptomycin, supplemented with10 μM of Y-27632 (ROCK inhibitor; Tocris Bioscience) and10 μM SB431542 (TGFBR1 inhibitor; Tocris Bioscience). To obtain a polarized epithelial monolayer, cells were pooled from 10 wells, washed in PBS and trypsinized before seeding onto a 75 mm, 0.4 μm pore polycarbonate membrane (Transwell, Corning) coated with Type IV collagen (34 μg/mL) in sterile cell culture water. CM was added to upper and lower wells, and maintained at 37°C, 5% CO_2_ for five days. Fresh CM was added every two days. Following confluency, media was replaced with cell differentiation media (DM) containing 5% CM in DMEM/F12 medium supplemented with 20% FBS, 2 mM L-glutamine and 10 μM Y-27632. Fresh DM was added daily for two days for monolayer differentiation prior to treatment with toxin.

### Spheroid propagation and processing

For spheroid cultures, cells suspended in Matrigel were seeded as 15 μL droplets into individual wells of a 48-well tissue culture plate and grown for 3 days in 50% CM. Spheroids were then diluted in 150 μL Matrigel (1:10 dilution from initial volume), re-seeded as 50 μL droplets in a 24-well tissue culture plate, and grown for 5 days in 50% CM. Finally, CM was replaced with cell differentiation media containing 5% CM and grown for 3 days. Mucin production was stimulated by the addition of *E. coli* heat-labile toxin LT [100 ng/ml]. Resulting spheroids were fixed with 4% paraformaldehyde (PFA) for 30 min at 37°C and 30 min at room temperature, washed 3x with PBS, and embedded in 5% Luria-agar prior to being processed as paraffin blocks for sectioning.

### Confocal microscopy

For EatA binding, spheroid sections were deparaffinized, hydrated, unmasked, and incubated with 10 μg/ml of EatAH134R or EatAH134R-DUF-his at 4°C overnight. Following incubation unbound EatA was washed off by rinsing the slides 3x with PBS. Muc2 mucin and EatA signals were detected with anti-Muc2 rabbit polyclonal (1:200, sc-15334, Santa Cruz) and anti-EatA mouse polyclonal (1:100, in-house) primary antibodies, followed by fluorescence tagged goat anti-rabbit IgG Alexafluor 488 (1:200, Invitrogen) and fluorescence tagged goat anti-mouse IgG Alexafluor 594 (1:200, Invitrogen) secondary antibodies, respectively. Cell nuclei were counterstained with DAPI (1:1000) and mounted using prolong gold antifade reagent (Invitrogen). Images were captured and analyzed on a Nikon C2 confocal microscope equipped with NIS-Elements AR 5.11.01 software (Nikon).

### Human intestinal explants and two-photon imaging

Freshly isolated sections of human small and large intestine were obtained at ileocolonic resection surgery from the Digestive Diseases Research Core Center at Washington University (DDRCC). The muscle layer was carefully removed and the tissues were supported on a 70 μm cell strainer mesh placed in a custom imaging chamber. The tissue was perfused from the serosal side with oxygenated DMEM throughout imaging. Wild type (jf2450) and *eatA* mutant (jf4738) GFP-expressing bacteria were grown in Luria broth containing ampicillin (100 μg/ml) overnight at 37°C, 200 rpm, and then diluted 1:100 and grown to OD_600_ of ~0.125 immediately prior to the experiment. Four hundred μl of undiluted culture was then mixed with 50 μl of a 1:100 dilution of 1.75 μm fluorescent microspheres (Polysciences 19392). Four hundred fifty μl of the bead/culture suspension was added to the epithelial surface of individual tissue sections. Images were acquired using a custom-built two photon microscope in the Washington University In-*vivo* Imaging Core (IVIC) equipped with an Olympus XLUMPLFLN 20x 1.0 N.A. water dipping objective. Fluorescence was excited using a Ti-Sapp laser (Chameleon Vision II, Choherent) tuned to 890 nm and emission collected using 495/540/570 nm dichroic filters (Semrock) and 3 head-on bi and multialkali PMTs (Hamamatsu). Slidebook digital microscopy software (Intelligent Imaging Innovations) was used for hardware control and acquisition of 2D timelapse recordings and z-stack images. Imaris 9.5 (Oxford Instruments) was used for data rendering and analysis.

### Metabolic labeling of enteroids

Enteroid monolayers were metabolically labeled overnight with media containing 50 μM GalNaz (N-azidoacetylgalactosamine tetraacylated, Jena Biosciences CLK-1086). After washing with fresh media, resulting azide-functionalized glycoconjugates were reacted with DBCO-Cy3 (Kerafast), and nuclei were stained with Hoescht 3342 (Sigma, B2261).

### Mucin purification

Human MUC2 mucin was recovered from culture supernatants of LS174T cells (ATCC CL-188) by ultrafiltration as previously described(21).

### Mucin migration assays

To evaluate bacterial migration through a mucin network, mucin was first harvested from human small intestinal enteroid monolayers. Approximately 48 h after differentiation, media was aspirated and mucin was carefully retrieved using a 1.8 cm wide cell scraper (Corning), leaving the cell monolayer undisturbed. 200 μL of the harvested mucin was dispensed onto 3 μm polycarbonate Transwell membranes in a 24-well plate. Five hundred μl of base media (Advanced DMEM/F12 containing 20% FBS and 2 mM L-glutamine) was added to the bottom wells. Fifty μl of bacterial suspension containing 10^5^ cfu was dispensed into the wells containing mucin, and plates were incubated at 37°C, 5% CO_2_, for 2h. Media was then collected from the bottom well, and serial dilutions were then plated on Luria agar with appropriate antibiotics. Purified EatA protein was added, where indicated, at a final concentration of 50μg/ml.

To evaluate the impact of EatA on migration through mucin to the epithelial surface, the upper chamber of monolayer cultures of small intestinal enteroids was inoculated by with ~10^5^ cfu of wild type or mutant bacteria. Plates were then incubated at 37°C, 5% CO_2_, and after 2 h media was gently aspirated, washed 3 times with pre-warmed media to remove unbound bacteria, and cells were then fixed with 4% paraformaldehyde for 30 min at room temperature. The fixed samples were then blocked with phosphate buffer solution (PBS) containing 1% bovine serum albumin (BSA) for 30 min, immunostained with rabbit anti-O78 antibody followed by cross absorbed goat anti-rabbit AlexaFluor 594-conjugated IgG (H+L) (ThermoFisher A-11072). Membranes with fluorescently labeled samples were excised and mounted onto glass slides with ProLong Gold (ThermoFisher Scientific P36930), and imaged with a Zeiss Axio Imager M2 Plus Wide Field Fluorescence Microscope, capturing 10 random fields per membrane to enumerate bacteria/field.

To evaluate the impact of EatA on kinetics of ETEC migration through mucin, equal numbers of GFP-expressing *eatA* mutant (jf4737) or wild type H10407 (jf2451) were loaded into adjacent wells of μ-channel slides (ibidi, vi 0.4) containing MUC2+ concentrated supernatants from LS174T cells. Slides were maintained at 37°C, 5% CO_2_, and 80% humidity in a stage top environmental chamber (ibidi). Confocal images were collected at 3 min intervals, at distances of 1.8, 4.5, 8.8, and 13.7 mm from the inoculation well, and mean fluorescence intensity was recorded over time to assess migration of GFP-expressing bacteria.

### Mucin integrity assays

To examine the impact of EatA and EatA mutants on mucin integrity we used a micro-scale rolling ball viscometer apparatus as described by Tang(45). Briefly, purified mucin treated with protease or controls were loaded into a 100 mm glass capillary tube (smurray G119/02) sealed at one end with modeling clay (Plastalina, Van Aken). 0.5 mm steel balls were introduced into the open end of the tube and maintained in place with a magnet. Tubes were maintained on a 20° incline, and the time required for the ball to traverse a distance of 80 mm following release of the magnet was recorded averaging a minimum of 4 technical replicates/experiment.

### Immunoblotting

MUC2 immunoblotting was performed as described previously(21). Briefly, after EatA treatment, samples were separated by discontinuous 4% tris/glycine SDS-PAGE, the gel was then reduced by agitation for 10 min in a 10 mM DTT solution in tris/glycine/isopropanol blotting buffer to improve transfer of the high-molecular mass MUC2, and then blotted onto nitrocellulose, blocked with 5% (w/v) milk in PBST, and probed with anti-MUC2 rabbit polyclonal (H-300, sc-15334; Santa Cruz) [1:2,000], followed by goat anti-rabbit-HRP conjugate sc-2030 Santa Cruz [1:5,000]. Bound antibody was visualized by chemiluminescence using Clarity ECL western blot substrate (Bio-Rad).

### MUC2 dot blotting

MUC2 dot blots were performed with supernatants from the apical compartment of small intestine enteroid monolayers grown as described above. Briefly, cells were grown in Transwells to confluence, differentiated for 2 days, then duplicate wells were treated overnight with 100ng/mL of native LT, mutant LT (E112K, mLT), or cholera toxin (CT). Supernatants were collected and spun down at 10,000 x g for 1 min to remove cellular debris. Serial dilutions (1:2 to 1:64) of the corresponding supernatants were performed in PBS. Serial dilutions (1:2 to 1:128) of concentrated LS174T supernatant containing MUC2 (described above) were used as positive controls. Two μl of each dilution was spotted onto a nitrocellulose membrane and allowed to dry, then blocked with 5% (w/v) milk in PBST, and probed with anti-MUC2 (clone F2, sc-515032, Santa Cruz) [1:500] followed by horse anti-mouse-HRP conjugate (7076, Cell Signaling) [1:1000]. Bound antibody was visualized by chemiluminescence using Clarity ECL western blot substrate (Bio-Rad).

### Glycan array screening

Glycan arrays were used as previously described(46) to examine the potential interaction of the EatA passenger domain with carbohydrates. Briefly, these glycan arrays which contained 737 separate features (including control spots) were fabricated, validated and analyzed as reported earlier (47–50). After blocking overnight at 4°C, using 3% BSA in PBS (w/v; 200 μl/well) slides were washed 6 times with PBST (PBS with 0.05% Tween-20, 200 μl/well). Biotinylated rEatA-6His, or biotinylated rEatA_p_-DUF∷His_8_ were diluted to final concentrations of 5 and 50 μg/ml in lectin buffer 20 mM Tris, 150 mM NaCl, 2mM MgCl_2_, 2mM CaCl_2_, pH 7.4) containing 3% BSA and 1% HSA (MilliporeSigma). In one pair of assays, 100 μL of each sample was incubated on the arrayfor 2 h (37°C, 100 rpm). The arrays were washed and incubated (1 h at 37 °C; 100 rpm) with Cy3-labeled streptavidin (MilliporeSigma, S6402) at a final concentration of 2 μg/mL in lectin buffer (1 h at 37 °C; 100 rpm). In a second pair of experiments, each sample was pre-complexed with streptavidin (10:1 protein:streptavidin), followed by incubation on the array for 2 h (37°C, 100 rpm). Slides were then washed 7 times in PBST (200 μL/well), immersed in wash buffer (5 min at room temperature), then centrifuged (5 min at 200 × *g*). Slide scanning was performed on an InnoScan 1100AL fluorescence scanner (Innopsys; Chicago, IL) at 5 μm resolution (Ex_532_/Em_575_) nm and data analyzed using GenePix Pro 7.0(47). A complete list of array components are available in supplemental dataset 1.

### Molecular cloning

To produced properly folded polyhistidine-tagged EatA passenger domain, the *eatA* gene was first amplified in two fragments using primer pairs jf110414.1/jf02215.5 and jf022515.6/jf022515.7 (table 3), yielding amplicons *Nco*I-*eatA*_5-258_8His-*Xho*I, and *XhoI*-*eatA*_259-4095_-*HindIII*, respectively. These fragments were sequentially introduced into the corresponding restriction sites on pBAD/myc-HisB (table 2) to produce pTV001 which encodes EatA bearing a polyhistidine tag at a permissive site between amino acids 86-87. Similarly, a plasmid was constructed to replace the region encoding the domain of unknown function corresponding to E541-S616 of EatA with an 8X polyhistidine tag. Briefly, primer sets jf110414.1/jf061615.3 and jf061615.2/jf022515.7 were used to generate amplicons *Nco*I-*eatA*_5-1620_9His-*XhoI* and *XhoI-eatA*_1849-4095_-*HindIII*, respectively. These fragments were then sequentially cloned into the corresponding sites on pBAD/myc-HisB, yielding pTW5126.

To generate the *eatA* complementation plasmid, pTW5016, full length *eatA* DNA sequence was amplified from H10407 with primer set jf081718.1 and jf081718.2. The PCR product was ligated to a low-copy plasmid vector pWSK29 (*BamH*I/*Sal*l) by Gibson assembly (New England Biolabs). Similarly, pTW5128 was constructed by amplifying the mutated *eatA* sequence from pTW5126 with primers jf071519.1 and jf071519.2. The amplicon was then ligated into *PstI-digested* pTW5016 by Gibson assembly.

The region corresponding to K535-P618 of EatA was amplified using primers jf070220.1/.2, and cloned by Gibson assembly to replace the mClover3 sequence of pSpyTag003-mClover3 which was amplified using primers jf070220.3/.4. DNA sequence of the resulting plasmid pJMF0820 confirmed placement of the region encoding EatA subdomain in-frame with 5’ SpyTag, and 3’ 6His tags.

Plasmids pTV001 and pTW5126 are deposited in addgene with accession numbers 130265 and 129732, respectively.

### Recombinant protein expression and antibody purification

To prepare untagged recombinant native (rEatA_p_) and inactivated (rEatA_p-H134R_), passenger domains proteins were expressed from strains jf1960 and jf1975 (supplemental table 1), and concentrated culture supernatants were then purified by anion exchange followed by size-exclusion chromatography, as previously described(21). The mutant passenger domain in which the domain of unknown function was replaced by a polyhistidine tag (rEatA_p_-DUF∷His_8_) was expressed from jf5132. Concentrated culture supernatant was then exchanged into PBS, and the secreted recombinant protein purified using immobilized metal affinity chromatography, as described(51)

### Production of SpyCatcher-EatAK535-P618 nanoparticles

BL21-CodonPlus (DE3)-RIPL transformed with pJMF0820 was grown overnight in Luria-Bertani (LB) media containing kanamycin 50 μg/ml at 37°C. The following morning saturated cultures were diluted 1:100 in 1 liter of fresh LB media supplemented with kanamycin and 0.8% glucose (w/v) and grown at 37°C, 200 rpm to an A_600_ of ~0.5. Cultures were then induced with isopropyl ß-D-1-thiogalactopyranoside (IPTG, 0.5 mM) and incubated at 30°C for and additional 4.5 h. Following centrifugation bacterial pellets were frozen at −80°C. Each of 4 pellets from 250 ml of culture was resuspended 30 ml of IMAC lysis buffer 75 mM sodium phosphate, pH 7.5, 500 mM NaCl, 1mM imidazole, cOmplete, mini, EDTA-free protease cocktail (Roche), 1 mg lysozyme, 1 mg DNAse, and 0.5 % Triton-x-100). Following sonication, and centrifugation at 20,000 x g for 20 min, polyhistidine tagged recombinant proteins were purified from the clarified lysates by immobilized metal affinity chromatography (IMAC) as previously described(35), yielding SpyTagged-EatA_K535-P618_.

His6-SpyCatcher-mi3 was produced as previously described(34). Briefly, after transformation of BL21-CodonPlus (DE30)-RIPL cells to kanamycin resistance transformants were grown overnight in Luria Bertani broth containing kanamycin (50 μg/ml), chloramphenicol (15 μg/ml), and streptomycin (10 μg/ml), then diluted 1:100 into 4 x 500 ml flasks with fresh media, and allowed to grow at 37°C to OD_600_ of ~ 0.7, then induced with 0.5 mM IPTG, overnight at room temperature, 250 rpm. Bacteria were harvested by centrifugation, and lysed in 30 ml of IMAC lysis buffer. His SpyCatcher-mi3 was then purified using affinity precipitation by the addition of nickle chloride to clarified lysates at a final concentration of 200 μM, and incubation at room temperature for 10 min. Precipitated protein was collected by centrifugation at 11,000 x g for 10 min and the resulting pure protein pellet redissolved in 10 ml of 50 mM Tris, pH 9, 10 mM EDTA. Polyclonal IgG from the sera of mice previously immunized with rEatAp(21), or normal mouse controls, was purified by protein G affinity chromatography.

### Antibody preparation

Antibodies were purified from the sera of twenty mice that were vaccinated intranasally three times, two weeks apart with 20 μg of recombinant EatA passenger domain (rEatAp) mixed with 1 μg of heat-labile toxin, as described previously(21). Two weeks after the final dose, mice were sacrificed and serum collected. The sera were screened for αEatAp IgG antibodies by ELISA, and ten sera with the highest αEatAp titres were pooled. To isolate antibodies specific to the EatA domain of unknown function from K535-P618, an affinity column was prepared by coupling 3.6 mg of the SpyTag-EatA_K535-P618_-His protein to AminoLink plus resin (Thermo) in 100 mM sodium citrate, 50 mM sodium carbonate, pH 10, at room temperature in a 10ml glass column for 4 h. The resin was then washed with 5 ml of PBS and 5 ml 50 mM sodium cyanoborohydride in PBS was added and allowed to react overnight at 4°C. The resin was then washed with 4 ml of quenching buffer (1M tris, pH 7.4). 2 ml of quenching buffer containing 50mM sodium cyanoborohydride was added and the suspension mixed at room temperature for 30 min. The resin was then washed with 10ml PBS prior to antibody capture.

Pooled αEatAp sera were diluted 1:1 with PBS, and mixed with the resin for 1 h at room temperature. The resin was then washed three times with 5 ml of PBS, and bound antibodies were then eluted in 10 ml 100 mM glycine, pH 2.7. 1 ml fractions of eluate were collected in tubes containing 50 μl of neutralisation buffer (1M Tris, pH 9). Fractions containing αDUF antibodies by ELISA were then pooled and dialysed against PBS.

### Toxin delivery assays

Enteroid-derived monolayer of polarized cells were obtained in a 24-well cell culture plate using the method described above. Cells cultures were treated with phosphodiesterase (PDE) inhibitors vardenafil (MilliporeSigma, Y0001647), cilostazol (MilliporeSigma, PHR1503), and rolipram (MilliporeSigma, R6520) 1 h prior to infections at a final concentration of 25 μM. The cells were then inoculated by dispensing 100 μl containing ~10^5^ cfu per to individual wells. The samples were incubated at 37°C in a 5% CO_2_ incubator for 2h, after which the media was replaced with fresh, prewarmed media containing PDE inhibitors, and incubated for 1 h. Cellular cAMP (Arbor Assays) levels were used as a readout for the efficiency of LT (heat-labile toxin) delivery as previously described (52)

### Oligopeptide cleavage

p-Nitroanilide substrate cleavage was carried out as previously described(18). Briefly, 1 mM N-Succinyl-Ala-Ala-Pro-Leu-p-nitroanilide (Sigma S8511) dissolved in 100 mM MOPS pH 7.3, 200 mM NaCl was digested at 37°C with 15 μg of enzyme in a final volume of 300 ul. The reaction was followed at 405 nm and initial rates expressed as mOD/min.

### Molecular modeling

The structure of the EatA passenger domain (GenBank accession AAO17297.1, amino acids 57-1062) was aligned with the corresponding region of SepA(33) using the Multialign Viewer extension(53) within UCSF Chimera(54) v1.13.1, developed by the Resource for Biocomputing, Visualization, and Informatics at the University of California, San Francisco, with support from NIH P41-GM103311. Theoretical homology models of the EatA passenger were generated against the SepA template Protein Data Bank entry 5J44 using the Chimera interface to Modeller (55).

### Ethics statement

Studies included here involving human tissues were approved by the Institutional Review Board at Washington University in Saint Louis School of Medicine. De-identified human small intestinal enteroids and fresh explants of human intestine were obtained from the DDRCC under approved protocols 201406083 and 201804112, respectively. Mouse studies were conducted under protocol 20-0438 approved by the Institutional Animal Care and Use Committee at Washington University in Saint Louis School of Medicine.

## Acknowledgements

JMF was supported by funding from the National Institute of Allergy and Infectious Diseases (NIAID) of the National Institutes of Health (NIH) R01 AI126887, R01 AI089894, U01 AI095473;, and the Department of Veterans Affairs (5I01BX001469-05). Research conducted by AS was also supported by National Institute of Allergy and Infectious Diseases of the National Institutes of Health under Award Number T32AI007172. The content is solely the responsibility of the authors and does not necessarily represent the official views of the National Institutes of Health, or the Department of Veterans Affairs.

BA and MP were supported in part by the Institute for Public Health Summer Scholars Program at Washington University, in Saint Louis. MJM was supported by funding from NIH/NIAID R01 AI077600. Intestinal biopsy specimens were obtained through the Biobank Core of the Digestive Disease Research Core Center (DDRCC) supported by NIH Washington University DDRCC Grant No. NIDDK P30 DK052574, and two-photon imaging experiments were conducted at the at Washinton University School of Medicine, *In vivo* Imaging Core (IVIC). SvdP, and GCH were supported by Swedish Research Council (Vetenskapsrådet, 2017-00958, 2020-02536).

